# The basolateral amygdala to lateral septum circuit is critical for regulating sociability in mice

**DOI:** 10.1101/2021.10.21.464669

**Authors:** Lionel A. Rodriguez, Sun-Hong Kim, Stephanie C. Page, Claudia V. Nguyen, Elizabeth A. Pattie, Henry L. Hallock, Jessica Valerino, Kristen R. Maynard, Andrew E. Jaffe, Keri Martinowich

**Affiliations:** Department of Neuroscience, Johns Hopkins School of Medicine, Baltimore, MD, 21205, USA; Lieber Institute for Brain Development, Johns Hopkins Medical Campus, Baltimore, MD, 21205, USA; Department of Genetic Medicine, McKusick-Nathans Institute of Genetic Medicine, Johns Hopkins University School of Medicine, Baltimore, MD, 21205, USA; Department of Psychiatry and Behavioral Sciences, Johns Hopkins School of Medicine, Baltimore, MD, 21205, USA; Center for Computational Biology, Johns Hopkins University, Baltimore, MD, 21205, USA; Department of Biostatistics, Johns Hopkins Bloomberg School of Public Health, Baltimore, MD, 21205, USA; Department of Mental Health, Johns Hopkins Bloomberg School of Public Health, Baltimore, MD, 21205, USA

**Author notes:** Equal Contribution (Lionel A. Rodriguez, Sun-Hong Kim). Correspondence: Keri Martinowich, Lieber Institute for Brain Development, 855 North Wolfe Street, Office #382, Baltimore, MD, 21205.

## Abstract

The lateral septum (LS) is a GABAergic region in the basal forebrain that is implicated in sociability. However, the neural circuits and cell signaling pathways that converge on the LS to mediate social behaviors aren’t well understood. Multiple lines of evidence suggest that brain-derived neurotrophic factor (BDNF) signaling through its receptor TrkB plays important roles in social behavior. While BDNF is not locally produced in LS, we demonstrate that nearly all GABAergic neurons in LS express TrkB. Local knock-down of TrkB expression from LS neurons decreased sociability and reduced recruitment of social novelty-induced neural activity. Since BDNF is not synthesized in LS, we evaluated which inputs to the LS could serve as potential BDNF sources for controlling sociability. By selectively ablating inputs to LS, we demonstrated that inputs from the basolateral amygdala (BLA), but not ventral CA1 (vCA1), regulate sociability. Moreover, depleting BDNF selectively in BLA-LS projection neurons phenocopied the decreased sociability observed following either local LS TrkB knockdown or ablation of BLA-LS inputs. These data support the hypothesis that BLA-LS projection neurons could serve as a critical source of BDNF for activating TrkB signaling in LS neurons to control sociability.

## INTRODUCTION

Social deficits are prevalent in many psychiatric and neurodevelopmental disorders, including autism [1,2] and schizophrenia [3]. While changes in social behavior manifest differently across these disorders, deficits in attention to socially salient stimuli and social recognition are well documented [3–5]. Mice are a highly social species that display robust responses to novel conspecifics, such as attention to, and investigation of, novel individuals [6,7]. The three-chamber social approach task, which measures the amount of time a mouse spends in close proximity with another mouse, was developed and validated to assess sociability, social recognition, and social preference [8]. Although rodent models cannot fully recapitulate the complexity of human social behavior, they are important for studying neural circuits that are linked to social behaviors observed in human disorders.

The lateral septum (LS) is a spatially complex, predominantly GABAergic region that extends across a significant portion of the rostral-caudal axis of the basal forebrain, and is subdivided into dorsal, intermediate and ventral subregions [9]. The LS is a potent modulator of social behaviors in both humans [10–13] and rodents [9,14] with documented roles in social saliency [15,16], social recognition [17–19], and social aggression [20,21]. The LS is innervated by many brain regions with established roles in social behaviors, including the basolateral amygdala (BLA) [22], ventral hippocampus CA1 area (vCA1) [23], ventral tegmental area [9,24], as well as the infralimbic and prelimbic cortices [9,25,26]. However, the inputs to the LS that mediate social recognition are not well understood, and the cell-signaling mechanisms in the LS that transmit this social information remain unclear.

Multiple lines of evidence suggest that the activity-dependent neurotrophin -brain-derived neurotrophic factor (BDNF) - is required for a variety of social behaviors [27–30]. Signaling through its cognate receptor tropomyosin kinase B (TrkB), BDNF regulates dendritic morphology, synapse formation and synaptic plasticity [31–33]. Importantly, BDNF plays a critical role in supporting GABAergic neuron function [34–38]. GABAergic neurons rarely synthesize BDNF, but robustly express TrkB [39,40]. GABAergic neurons rely on BDNF released from other cell types to activate TrkB signaling, which is critical for their maturation and physiological function [37,41–43]. In line with its GABAergic composition, the LS is largely devoid of *Bdnf* transcript expression, but many neurons innervating the LS densely express BDNF [44,45]. Previous research has shown that BDNF-expressing catecholaminergic projections to the LS influence the morphology and gene expression of a subpopulation of calbindin-expressing LS neurons [46]. However, whether BDNF-TrkB signaling in the LS is important for the social behaviors mediated by the LS has not been investigated.

We hypothesized that 1) TrkB signaling in LS GABAergic neurons is important for regulating rodent social behavior, and that 2) neuronal projections from BDNF-rich limbic regions serve as a source of BDNF to activate TrkB signaling in LS to control this behavior. Key candidate regions included the BLA and vCA1 because both regions highly express BDNF [44], send projections to the LS [20,22,26], and have established roles in regulating social novelty [9,16,47] and recognition [23,48,49]. Indeed, our data demonstrate that local TrkB knockdown in the LS is critical for social novelty and recognition behavior in mice. Moreover, we identify BLA-LS projections as critical for these same behaviors, and show that this behavior depends on BDNF expression within these projection neurons. In summary, the data demonstrate a vital role for LS TrkB signaling in social novelty recognition, and support the possibility that BLA-LS neurons serve as a source of BDNF for activating LS TrkB receptors to control this behavior.

## MATERIALS AND METHODS

### Animals

Wild-type mice (C57BL/6J; stock # 000664) and mice carrying a *loxP*-flanked *Bdnf* allele (Bdnf^tm3Jae^; stock # 004339, referenced in text as BDNF^*fl/fl*^) were purchased from The Jackson Laboratory. Mice carrying a *loxP*-flanked TrkB allele (strain fB/fB, referenced in text as TrkB^*fl/fl*^ [50,51]) were used for some experiments. TrkB^*fl/fl*^ mice were maintained on a C57BL/6J background. Adult male mice were housed in a temperature-controlled environment with a 12:12 light/dark cycle and *ad libitum* access to food and water. All experimental animal procedures were approved by the Sobran Biosciences Institutional Animal Care and Use Committee.

### Behavior

The three-chamber social interaction test arena consisted of three adjacent chambers separated by two clear plastic dividers, and connected by two open doorways. Mice were habituated for 3 consecutive days (10 min per day) prior to testing days. Mice were transferred to the testing room 1 h prior to testing. The test consisted of 5 min of habituation and two 10 min sessions (trial 1 and 2) with a 5 min inter-trial-interval. The subject mouse was kept in the center chamber during habituation. In the first 10 min session (trial 1), the subject mouse was allowed to freely investigate the three chambers. A novel mouse (stranger 1) was placed under an inverted metal cup in one side chamber, an identical empty cup was placed in the other side chamber. After the trial 1 session, the doorways were closed and the subject mouse was kept in the center chamber for 5 min. In the second session (trial 2), another novel object mouse (stranger 2) was put in the empty cup and the subject mouse freely investigated the three chambers. Age-matched adult C57BL/6J male mice were used as object mice in this study, and housed under the same conditions as the subject mice. Each object novel/stranger mouse was used once a day in the three-chamber social interaction test. Time spent sniffing each cup was manually scored as interaction time in a blind manner using the Stopwatch+ program developed by the Center for Behavioral Neuroscience (cbn-atl.org) at Emory University [52,53].

For full description of methods, see Supplementary information.

## RESULTS

### Local knockdown of LS TrkB expression

We used RNAscope single-molecule fluorescent *in-situ* hybridization to quantify expression of the gene encoding TrkB (*Ntrk2* probe) in GABAergic neurons (*Gad1* probe) across the rostral-caudal axis of the LS (Fig. 1A,B). These data revealed that in wild-type mice, nearly all LS GABAergic neurons (85-90%) expressed *Ntrk2*, a pattern that was consistent across the rostral-caudal axis (Fig. 1B). Next, we designed a genetic manipulation strategy to knockdown expression of TrkB in the LS. Specifically, we bilaterally injected a cre recombinase-expressing virus (AAV5-EF1a:Cre) into the LS of mice carrying a floxed TrkB allele (TrkB^*fl/fl*^ mice) to knockdown TrkB selectively in LS (Fig 1C). As controls, we generated mice that retained intact TrkB expression (TrkB^*fl/fl*^ mice injected into the LS with a non-Cre expressing virus; AAV5-EF1a:EYFP) (Fig. 1C). Four weeks following injection of the viruses, we micro-dissected the LS and frontal cortex from each mouse, and quantified protein expression of both full-length and truncated TrkB in individual mice (Fig. 1D-F). Expression of cre recombinase in LS of TrkB^*fl/fl*^ mice resulted in no change in the frontal cortex, but ∼86% knockdown of full-length TrkB expression in LS (Fig. 1D,E). Original characterization of this TrkB floxed allele (TrkB^*fl/fl*^) crossed with the Dlx5/6 cre driver line caused complete loss of full-length TrkB expression, but also partial loss of truncated expression [51]. In contrast, we observed no change in truncated TrkB expression across experimental groups (Fig. 1D,F). This difference may be accounted for by differences in cell composition in the brain regions studied (striatum versus LS) and differences in method of cre delivery (Dlx5/6 Cre-driver mouse versus local delivery of AAV).

**Figure 1.**
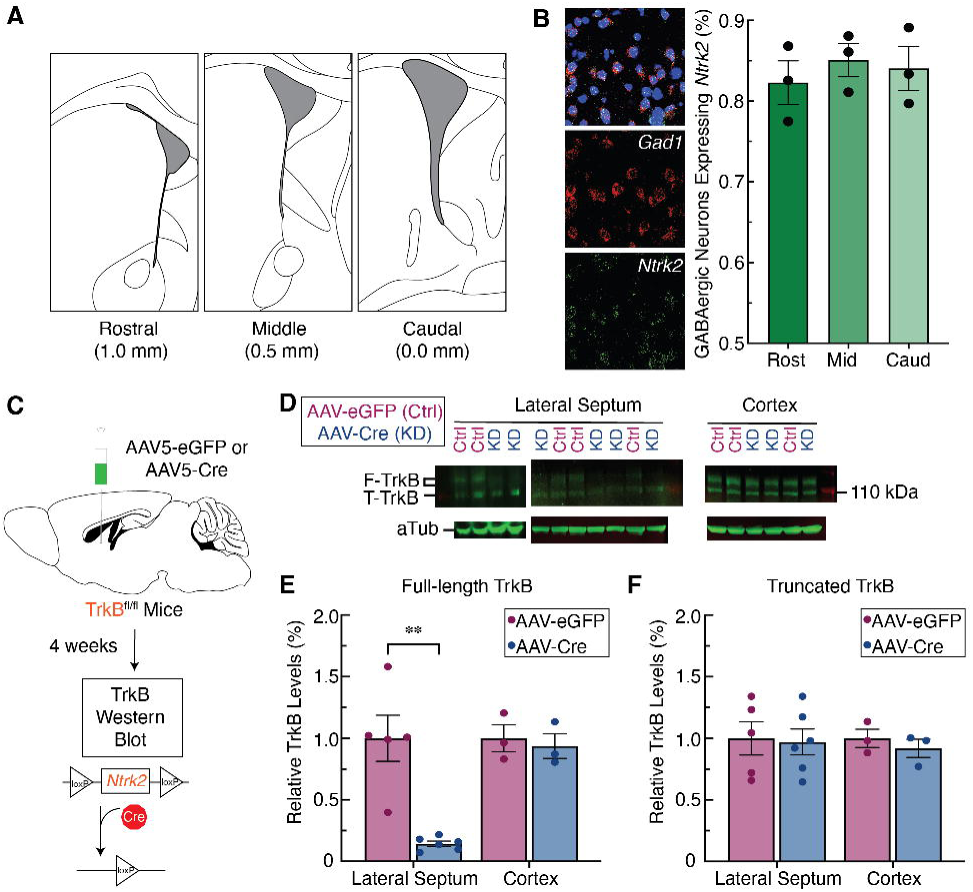
Tropomyosin receptor kinase B (TrkB) is highly expressed in the lateral septum (LS) and can be effectively depleted virally. (A) Illustration depicting the anatomical boundaries across the rostral-caudal axis of the LS. (B) Fluorescent *in-situ* hybridization in sections of mouse LS (n=4) demonstrates *Gad1*-positive cells highly express *Ntrk2* throughout the rostral-caudal divisions. (C) Viral strategy using Cre-induced TrkB knockdown in the LS of TrkB^fl/fl^ mice to quantify extent of TrkB knockdown quantification. (D) Western blot images for full-length TrkB (F-TrkB), truncated TrkB (T-TrkB), and α-Tubulin (aTub). (E) Relative expression levels of F-TrkB protein were significantly lower in the LS of the experimental group (AAV-Cre in LS of TrkB^fl/fl^ mice, n=5) compared to the control group (AAV-eGFP in LS of TrkB^fl/fl^ mice,n=6)(t_9_ = 5.015, p = 0.0014). (F) Relative expression levels for T-TrkB protein were similar between experimental and control mice.

### Knockdown of LS TrkB abolishes social novelty discrimination

We next used this local knockdown strategy to investigate the necessity of TrkB signaling in social novelty and sociability using the three chamber social interaction task (Fig 2A,B). This task uses a three-chambered box to quantify the time a mouse spends examining socially novel individuals. In trial 1, the mouse is exposed to a novel mouse (stranger mouse 1) in one of the outer chambers while the other outer chamber remains empty. In trial 2, stranger mouse 1 remains in the same chamber (now the familiar mouse) while a new mouse (stranger mouse 2) is placed in the other chamber.

**Figure 2.**
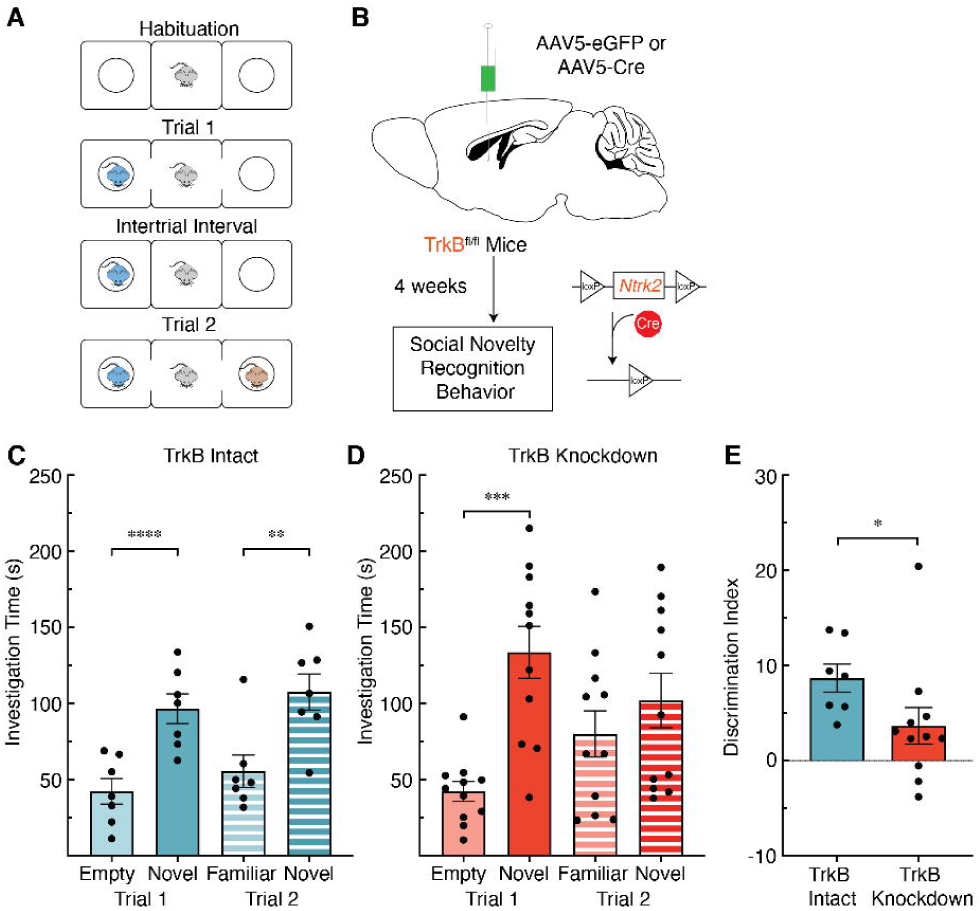
Tropomyosin receptor kinase B (TrkB) expression in the lateral septum (LS) is required for social novelty recognition behavior in mice. (A) Description of three chamber social interaction experiment. (B) Viral strategy using Cre-induced knockdown of TrkB expression in the LS of TrkB^fl/fl^ mice prior to social novelty recognition. (C) Mice with intact TrkB expression in the LS (n=7) display intact social novelty recognition behavior. In both trial 1 (t_6_ = 10.99, p < 0.0001) and trial 2 (t_6_ = 5.921, p = 0.001) control mice spend more time investigating the socially novel mouse. (D) TrkB knockdown in LS (n=11) abolishes the social novelty phenotype observed in trial 2 (t_10_ =1.898, p = 0.088), but not trial 1(t_10_ = 5.615, p = 0.0002). (E) Mice with intact TrkB expression in the LS show better social discrimination between the socially novel and familiar mice in Trial 2 when compared to mice with TrkB knockdown in the LS (U_11,7_ = 12, p = 0.0154).

Control mice with intact TrkB expression showed expected social novelty behavior, spending more time investigating the novel mouse in trial 1 and trial 2 (Fig. 2C). However, while mice with LS TrkB knockdown showed expected social novelty behavior in trial 1, these mice spent equal time with the familiar and novel mice in trial 2 (Fig. 2D). To evaluate differences in the ability of control and LS TrkB knockdown mice to discriminate between individuals we generated a discrimination index for trial 2, which is a percentage of the time spent with the novel individual minus the time spent with the familiar individual over the total trial time. The decreased discrimination index in LS TrkB knockdown mice suggests that loss of intact full-length TrkB may impact the ability to discriminate between individual mice, or increase sociability (Fig. 2E).

To rule out that the impairment in attention to social novelty discrimination was due to olfactory impairment, both controls and mice with LS TrkB knockdown were subjected to a four trial odor discrimination task (Fig. S1A). LS TrkB knockdown did not impair odor discrimination (Fig. S1B). We also assessed the necessity of TrkB signaling in LS for a number of fear-and anxiety-related behaviors that are known to engage LS circuitry. LS TrkB knockdown did not significantly decrease anxiety-like behavior in the elevated plus maze (Fig. S1C), nor did it impact the ability of LS TrkB knockdown mice to acquire, retrieve or extinguish fear memories (Fig. S1D). These data support the hypothesis that the deficits observed following knockdown of LS TrkB are specific to social behavior.

### LS TrkB knockdown abolishes socially induced c-Fos Expression

To map neural activity patterns in LS cells following exposure to novel social stimuli, we performed fluorescent immunohistochemistry for c-Fos across the rostral-caudal divide of the LS (Fig. 3A) in wild-type mice exposed to a social stimulus (a novel male mouse) compared to a non-social stimulus (a novel object)(Fig. 3B). c-Fos quantification was performed using the open-source biological imaging software Fiji (for detailed description of methods, see Supplementary Information and Fig, S2A). In the middle portion of the LS, significantly more cells expressed c-Fos in response to social stimuli compared to the non-social stimulus (Fig. 3C). This effect mapped most strongly to the intermediate and ventral subregions of the LS (Fig. 3D). The number of c-Fos expressing cells in all other subregions within the rostral (Fig. S2B) and caudal (Fig. S2C) portions of the LS remained relatively consistent across the social and non-social conditions suggesting that these effects on cell recruitment in the LS are restricted to the middle portion of the LS.

**Figure 3.**
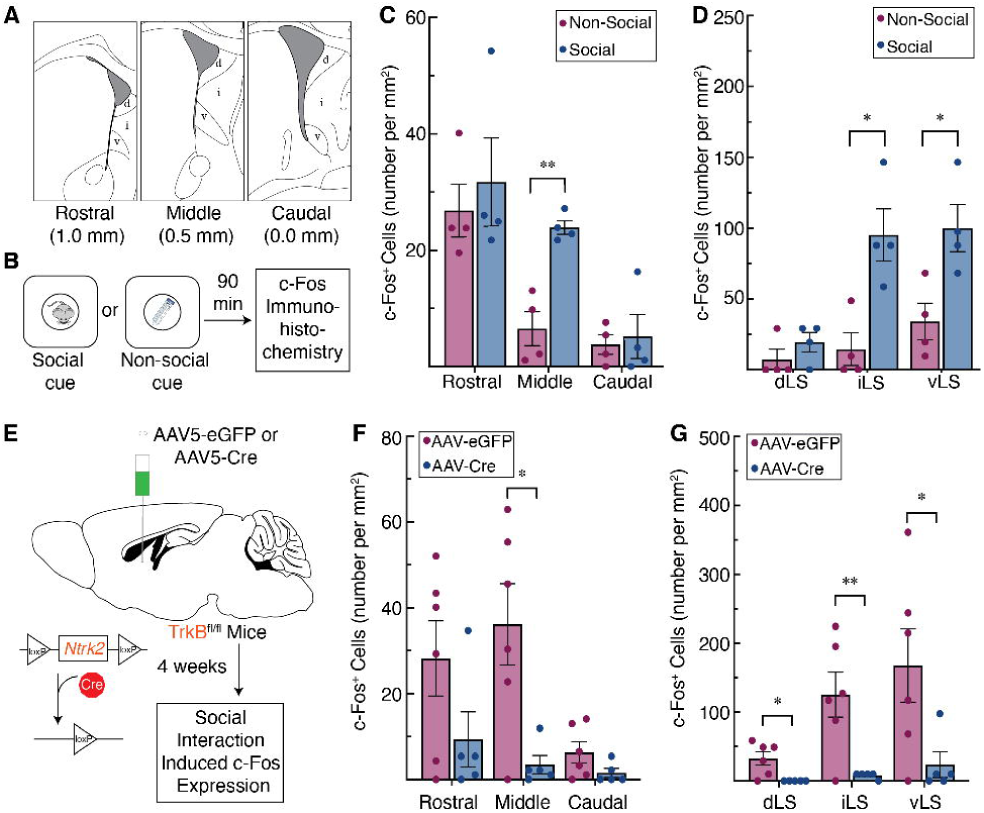
Tropomyosin receptor kinaseB (TrkB) expression in the lateral septum (LS) is required for social novelty induced c-Fos expression. (A) Illustration depicting the anatomical boundaries across the rostral-caudal axis of the LS, denoting the dorsal (d), intermediate (i), and ventral (v) subregions of the LS. (B) Description of social or non-social cues used to induce c-Fos expression. (C) Induction of c-Fos is significantly increased by social cues in the middle portion of the LS (t_6_ = 5.543, p = 0.004), (D) an effect seen specifically in the intermediate (t_6_ = 3.697, p = 0.030) and ventral subdivisions (t_6_ = 3.131, p = 0.040) of the middle LS. (E) Viral strategy using Cre-induced TrkB knockdown in the LS of TrkB^fl/fl^ mice prior to examining social interaction induced c-Fos expression. (F) Knockdown of TrkB in the LS significantly decreases cell recruitment in the middle portion of the LS following social stimuli (t_9_ = 3.069, p = 0.0396), (G) an effect seen across the dorsal (t_9_ = 3.049, p = 0.029), intermediate (t_9_ = 3.254, p = 0.029), and ventral (t_9_ = 2.342, p = 0.044) subdivisions of the middle LS.

To investigate whether knockdown of TrkB would abolish LS-responsivity to socially novel stimuli, we generated an additional cohort of LS TrkB knockdown and control mice using the same viral strategy described above (Fig. 3E). Compared to controls, we detected significantly fewer cells expressing c-Fos in response to novel social stimuli in the middle portion of the LS in LS TrkB knockdown mice (Fig. 3F). All subregions (dorsal, intermediate, and ventral) within the middle portion of the LS displayed blunted c-Fos expression in response to a socially novel individual (Fig. 3G). However, LS TrkB knockdown had no effect on c-Fos expression induced in response to socially novel individuals within the rostral (Fig. S3B) and caudal (Fig. S3C) portions of the LS compared to controls.

### BLA projections, but not vCA1 projections, to the LS are necessary for social novelty discrimination

While the LS is critical for controlling responses to social novelty [16–19], the brain regions that transmit information to the LS for proper execution of social novelty discrimination have not been identified. We noted the BLA and the vCA1 as candidate regions because both have established roles in regulating social novelty [9,16,47] and recognition [23,48,49], and send projections to the LS [22,26]. To determine the importance of projections from the BLA or vCA1 to the LS in controlling social novelty behavior, we used a diphtheria toxin A (DtA)-mediated viral circuit elimination strategy. Specifically, we bilaterally injected two viruses with retrograde tropism (AAVrg-CB7.CI:EGFP and AAVrg-EF1a:mCherry-IRES-cre) into the LS, causing expression of eGFP, mCherry, and Cre in cells projecting to LS (Fig. 4A). In our regions of interest (separate cohorts for BLA and vCA1) we also bilaterally injected a virus (AAV5-EF1a:mCherry-FLEX-dtA) expressing mCherry and cre-dependent DtA (Fig. 4A). In this scenario, cell bodies in the BLA or vCA1 that also sent projections to the LS expressed cre-recombinase, and hence cre-mediated expression of DtA in the BLA-LS or vCA1-LS neuronal projections. This manipulation caused elimination of the eGFP signal in those ablated projections, but retention of the mCherry signal in non-projecting cells (Fig. 4B). Control mice with an intact BLA-LS circuit showed expected social novelty behavior, spending more time investigating the novel mouse in trial 1 and trial 2 in the three chamber social recognition task (Fig 4C). Mice with ablated BLA-LS projections displayed expected social novelty behavior in trial 1, but in trial 2 these mice spent equal time with the familiar and novel mice (Fig. 4D). In addition, mice with intact BLA-LS projection neurons discriminated better than mice with ablated BLA-LS projection neurons according to their relative percentage of time spent with the novel versus familiar mouse in trial 2 (Fig. 4E). Both control mice with an intact vCA1-LS circuit (Fig. 4F) and a vCA1-LS ablated circuit (Fig. 4G) showed the expected social novelty behavior in trial 1 and trial 2. Neither group differed in ability to discriminate between the novel and familiar mice in trial 2 (Fig. 4H). These data support a role for BLA-LS neurons in regulating social novelty and sociability.

**Figure 4.**
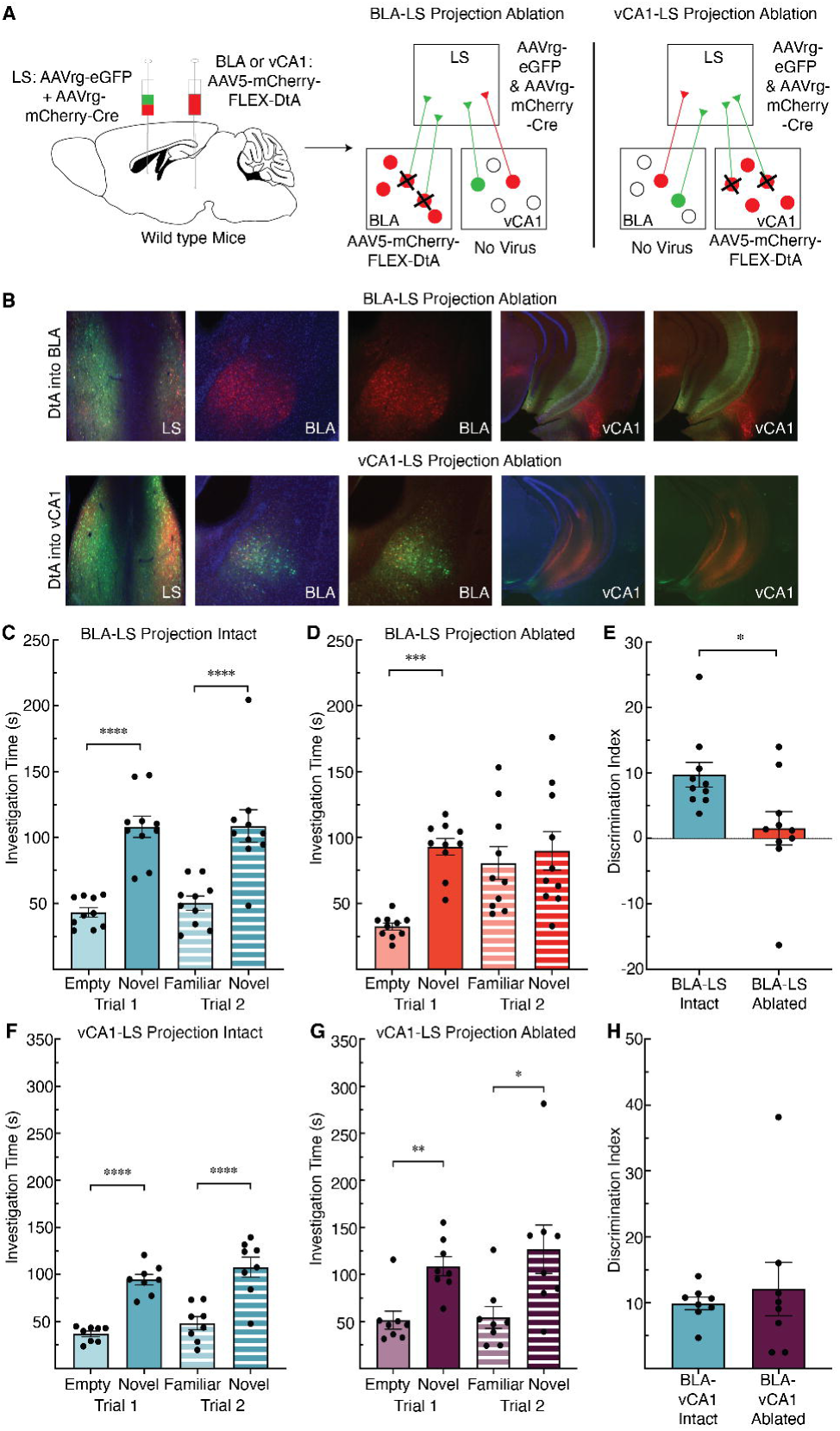
Ablating inputs to the lateral septum (LS) from basolateral amygdala (BLA), but not ventral CA1 (vCA1), abolishes social novelty recognition behavior in mice. (A) Cartoon of viral strategy used to selectively eliminate inputs to the LS from either the BLA or vCA1 utilizing retrograde labeling and Cre-mediated expression of diphtheria toxin A (DtA) (left panel). Schematics depict ablation in cohort where BLA-LS projections are targeted (middle panel), and ablation in cohort where vCA1-LS projections are targeted. (B) Representative images show that in the cohort where DtA is expressed in BLA (top row), BLA-LS projections are ablated, while vCA1-LS projections are intact, while in the cohort where DtA is expressed in vCA1 (bottom row), vCA1-LS projections are ablated while BLA-LS projections remain intact. (C) Mice with the BLA-LS circuit intact (n=10) display normal social novelty recognition behavior. In both trial 1 (t_9_ = 7.808, p < 0.0001) and trial 2 (t_9_ = 5.17, p = 0.0006) they spend more time investigating the socially novel mouse. (D) Ablation of the BLA-LS circuit (n=10) abolishes the social novelty phenotype observed in trial 2 (t_9_ = 0.6048, p = 0.56), but not trial 1(t_9_ = 12.74, p < 0.0001). (E) Mice with an intact BLA-LS circuit show better social discrimination between the socially novel and familiar mice in Trial 2 than mice with an ablated BLA-LS circuit (U_10,10_ = 17, p = 0.0115). (F) Mice with the vCA1-LS circuit intact (n=8) display normal social novelty recognition behavior. In both trial 1 (t_7_ = 8.449, p < 0.0001) and trial 2 (t_7_ = 10.5, p < 0.0001) they spend more time investigating the socially novel mouse. (G) Ablation of the vCA1-LS circuit (n=8) does not alter social novelty behavior in trial 1 (t_7_ = 5.235, p = 0.0012) or trial 2 (t_7_ = 2.979, p = 0.0206). (H) Mice with an intact vCA1-LS circuit and mice with an ablated vCA1-LS circuit show no difference in social discrimination (U_9,9_ = 30, p = 0.8527).

### Expression of BDNF within BLA projections to the LS is necessary for social novelty discrimination

Neurons in the BLA strongly express BDNF [44], and the BLA sends projections to the LS [22]. Hence, BLA-LS neurons are a strong candidate source for providing BDNF to LS TrkB-expressing neurons that regulate social novelty. To assess whether BDNF expression is necessary in BLA-LS projections to control social novelty behavior, we used a viral strategy combining the Flp recombinase (FlpO) and cre recombinase systems in mice carrying a floxed *Bdnf* allele (BDNF^*fl/fl*^). This strategy allowed us to eliminate BDNF selectively from BLA-LS projections (Fig. 5A). We bilaterally injected a retrograde virus in the LS (AAVrg-EF1a:Flpo) of BDNF^*fl/fl*^ mice, causing FlpO expression in all neuronal inputs to the LS. We simultaneously injected a virus (AAV8-EF1a:fDIO-mCherry-P2A-Cre) expressing Flp-dependent mCherry and cre-recombinase in the BLA. Control mice were injected with a virus (AAV8-EF1a:fDIO-mCherry) into the BLA that expressed Flp-dependent mCherry. In the experimental group, BLA neurons projecting to the LS express cre-recombinase in a FlpO-dependent manner. Expression of cre leads to excision of the floxed BDNF allele and Cre-dependent expression of mCherry (Fig. 5B). Control mice with intact BDNF expression in the BLA-LS circuit showed expected social novelty behavior, spending more time investigating the novel mouse in trial 1 and trial 2 in the three chamber social recognition task (Fig. 5C). Mice with knockdown of BDNF expression in BLA-LS projection neurons showed expected social novelty behavior in trial 1, but in trial 2 these mice spent equal time with the familiar and novel mice (Fig. 5D). Moreover, mice with decreased BDNF expression in BLA-LS projection neurons display poorer discrimination between individuals compared to animals with intact BDNF expression in BLA-LS projection neurons according to their relative percentage of time spent with the novel versus familiar mouse in trial 2 (Fig. 5E). In summary, BDNF expression in BLA-LS neurons is critical for social novelty discrimination.

**Figure 5.**
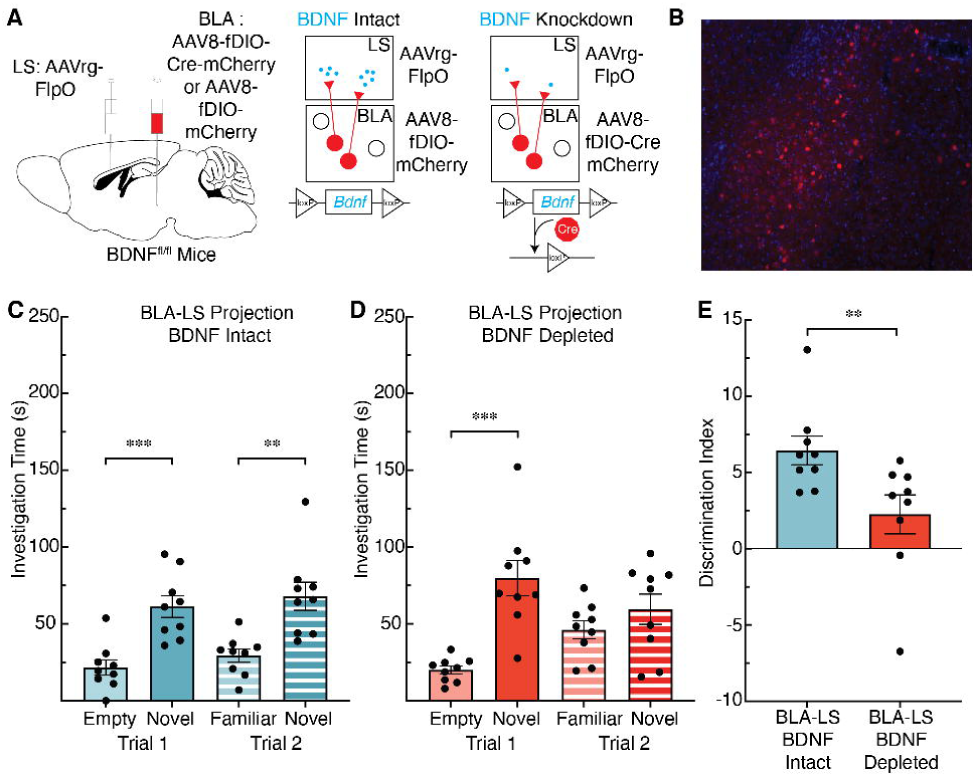
Knockdown of brain-derived neurotrophic factor (BDNF) in basolateral amygdala (BLA) inputs to the lateral septum (LS) abolishes social novelty recognition behavior in mice. (A) Schematic of viral strategy used to deplete BDNF expression in BLA-LS projections in BDNF^fl/fl^ mice. (B) Representative images of viral expression of mCherry within the BLA. (C) Mice with intact BDNF expression in BLA-LS projections (n=9) display normal social novelty recognition behavior. In both trial 1 (t_8_ = 5.73, p = 0.0004) and trial 2 (t_8_ = 6.842, p = 0.0001) they spend more time investigating the socially novel mouse. (D) BDNF depletion in BLA-LS projections (n=9 mice) abolishes the social novelty phenotype observed in trial 2 (t_8_ = 1.763, p = 0.116), but not trial 1(t_8_ = 5.456, p = 0.0006). (E) Mice with intact BDNF expression in BLA-LS projections display better social discrimination between the socially novel and familiar mice in trial 2 than mice with BDNF knockdown in BLA-LS projections (U_9,9_ = 9, p = 0.004).

## DISCUSSION

### BLA projections to the LS control sociability

The BLA is critical for valence processing in a variety of contexts [54,55], a function that may extend to processing social stimuli [56]. The BLA is responsive to socially novel conspecifics [16,47], and controls social discrimination [49], but which regions the BLA communicates with to regulate these behaviors is not well established. By demonstrating that ablating neuronal projections from the BLA to the LS abolishes social novelty behavior in the three chamber task, our data provide evidence that the BLA-LS circuit controls sociability. Because BLA neurons can collateralize [57], it is important to consider that ablating these BLA neurons may partially block transmission of social information to additional regions besides the LS. In addition to the BLA, vCA1 has been implicated in controlling responses to social novelty [16] and regulating social memories [23,48], and LS projections from adjacent subregions of the hippocampus, including dorsal CA2 (dCA2), are critical for controlling social aggression [20]. Surprisingly, ablating projections from vCA1 to the LS did not alter social discrimination behavior. While dCA2 projections to vCA1 are critical for social recognition [23], this social information may not be transmitted from ventral hippocampal circuitry to the LS to control social novelty and sociability. Further research would be necessary to determine if other portions of the ventral hippocampus, such as dCA2, regulate social novelty and recognition, in addition to aggression. Interestingly, activation versus inhibition of BLA projections to the ventral hippocampus bi-directionally controls time spent socializing [56], which may point to the BLA as a central transmitter of social information to limbic circuitry.

### TrkB signaling in the LS critically regulates sociability

The LS is a central hub for processing social behavior, and controls both social salience [15,16] and social recognition [17–19], but little is known about the cellular and molecular mechanisms by which social information is relayed to the LS. Knocking down TrkB expression in the LS abolished social novelty behavior in the three-chamber social interaction task, suggesting that TrkB signaling is necessary for discriminating socially novel conspecifics from those that are socially familiar. Importantly, TrkB knockdown did not impair olfactory discrimination, anxiety-like behavior, or fear learning, highlighting the specificity of LS TrkB signaling in controlling the response to social novelty.

The LS is spatially complex, containing dorsal, ventral, and intermediate subregions that span its rostral-caudal axis. Importantly, these subregions have functional distinctions, for example the dorsal and ventral LS control different aspects of social aggression in both males [23] and females [58]. While elevated c-Fos expression in the LS in response to socially novel stimuli is documented [15,16], how this activity maps to the spatial topography of the LS has not been rigorously explored. Using c-Fos neural activity mapping, we demonstrated that the intermediate and ventral subregions of the middle portion of the LS are the socially responsive loci. Previous results suggested that social novelty-induced activity in the LS was concentrated more rostrally than caudally [15], which is consistent with our results. In addition, we demonstrated that TrkB expression in the LS is necessary for LS responsiveness to social stimuli. Specifically, knockdown of LS TrkB expression blunts the recruitment of neural activity in the intermediate and ventral subregions of the middle portion of the LS in response to social stimuli.

### BDNF expression in BLA-LS neurons is critical for social novelty behavior

The BLA controls a variety of social behaviors in rodents, non-human primates [59] and humans [60–62]. While the cellular and molecular mechanisms that control these behaviors are complex, the prominent role of BDNF-TrkB signaling in many critical components of synaptic function renders it likely that this pathway is involved. Supporting this notion, rare genetic disorders that feature BDNF haploinsufficiency as well as common genetic variation in the human *BDNF* locus are linked with multiple components of social functioning, including aggression [63–65]. In addition to BDNF, a recent genome-wide association study of anxiety, which included human subjects with social anxiety, identified a single nucleotide polymorphism in *NTRK2*, the gene encoding the TrkB receptor, as contributing to genetic risk [66].

Our data demonstrate that TrkB expression in the LS is critical for social novelty and recognition behavior (Fig. 1D-G), but because BDNF is not synthesized in the LS, the source of BDNF for activating LS TrkB receptors is unclear [44]. BDNF is however highly expressed in BLA neurons [44], and given our data showing that BLA-LS projection neurons are critical for this behavior, we reasoned that BDNF produced in BLA-LS projection neurons could be a potential source for activating LS TrkB receptors to control this behavior. Supporting this possibility, knocking down BDNF selectively in BLA-LS projection neurons phenocopied the behavioral deficits caused by local TrkB knockdown in the LS. While our data support the hypothesis that release of BDNF from BLA-LS neurons into the LS engages local TrkB receptors to control social behavior, additional experiments would be needed to causally prove this link. In our experiments, BDNF expression in BLA-LS neurons is chronically depleted, which could lead to longer-term structural and physiological changes to BLA-LS projections due to the loss of neurotrophic support, including from decreased post-synaptic release of BDNF into the BLA itself. Such weakening of synaptic connections could lead to overall changes in the circuitry that could cause changes in social behavior. Moreover, while our data suggests that loss of TrkB expression in the LS gates the region’s response to social stimuli, further work with acute manipulations of TrkB receptors would be critical to understand the kinetics of how LS BDNF-TrkB signaling controls social behaviors. This is important since the chronic effects of TrkB knockdown in LS in our experiments could impact the physiological function of TrkB-expressing inhibitory neurons and alter the strength of inhibitory synapses. This in turn could impact connectivity within the LS, or inputs to the LS, contributing to the observed impairments in social behaviors.

### Ethological Considerations on Behavior

A limitation of these studies is that they were conducted only in male mice. While male and female mice show substantial overlap across social behaviors, there are documented differences, including establishing social hierarchies [67,68], neural coding of social behaviors [69], and processing social recognition [70]. Hence, direct comparisons of males and females in social novelty and recognition paradigms poses challenges in behavioral interpretation. Despite these challenges, there are several reasons why future studies should examine a potential role for BDNF-TrkB signaling in BLA-LS circuitry in the context of sex-specific social behaviors. First, prevalence rates [71], as well as genetic risk, for disorders that often feature social deficits are not equal across sex and gender in humans [72–76]. Second, BDNF-TrkB signaling and expression is significantly impacted by sex and hormones [77–79]. Conducting these studies, however, will require taking into consideration specific ethological aspects of female social behaviors. For example, female mice establish different social hierarchies depending on the presence of male mice [68]. This manifests in territoriality and aggression in dominant female mice, a behavior normally only seen in postpartum dams in defense of their pups [80]. Conversely, male mice naturally display this territoriality and aggression toward male mice [68,81], but seldom display these behaviors in male-female interactions [68]. Despite the difficulties in making direct comparisons between male and female mice in studies of social behaviors, well-designed experiments can be implemented to study sex-specific differences in social novelty and recognition. For example, studying female social novelty in postpartum mice or dominant female mice raised in complex social structures will elucidate whether the circuits that enable normal male responses to social novelty are also necessary for controlling female social novelty.

## Conclusions

Collectively, the data demonstrate a critical role for TrkB signaling in LS neurons in controlling social novelty recognition, and support the hypothesis that the BLA provides a critical source of BDNF for activating TrkB receptors in LS neurons to control this behavior. The data provide novel information about the broader role of signaling in amygdala-septal circuits, and establish a foundation for future studies that delineate how molecular and cellular signaling in these circuits impacts social functioning.

## ACKNOWLEDGEMENTS

We thank members of the Martinowich and Maynard laboratories for helpful comments and suggestions.

## AUTHOR CONTRIBUTIONS

LAR: Formal analysis, Investigation, Writing - Original Draft, Visualization

SHK: Conceptualization, Methodology, Formal analysis, Investigation, Writing - Review & Editing

SCP: Investigation, Resources

CVN: Investigation

EAP: Investigation

HLH: Investigation

JV: Investigation

KRM: Writing - Review & Editing, Methodology

AEJ: Formal analysis, Resources

KM: Supervision, Writing - Original Draft, Project administration, Funding acquisition

## REFERENCES

1. Chevallier C, Kohls G, Troiani V, Brodkin ES, Schultz RT. The social motivation theory of autism. Trends Cogn Sci (Regul Ed). 2012;16:231–239.

2. Happé F, Ronald A. The “fractionable autism triad”: a review of evidence from behavioural, genetic, cognitive and neural research. Neuropsychol Rev. 2008;18:287–304.

3. Marwick K, Hall J. Social cognition in schizophrenia: a review of face processing. Br Med Bull. 2008;88:43–58.

4. Mier D, Kirsch P. Social-Cognitive Deficits in Schizophrenia. Curr Top Behav Neurosci. 2017;30:397–409.

5. Weigelt S, Koldewyn K, Kanwisher N. Face identity recognition in autism spectrum disorders: a review of behavioral studies. Neurosci Biobehav Rev. 2012;36:1060–1084.

6. Pobbe RLH, Pearson BL, Defensor EB, Bolivar VJ, Blanchard DC, Blanchard RJ. Expression of social behaviors of C57BL/6J versus BTBR inbred mouse strains in the visible burrow system. Behav Brain Res. 2010;214:443–449.

7. Moy SS, Nadler JJ, Perez A, Barbaro RP, Johns JM, Magnuson TR, et al. Sociability and preference for social novelty in five inbred strains: an approach to assess autistic-like behavior in mice. Genes Brain Behav. 2004;3:287–302.

8. Lo S-C, Scearce-Levie K, Sheng M. Characterization of social behaviors in caspase-3 deficient mice. Sci Rep. 2016;6:18335.

9. Sheehan TP, Chambers RA, Russell DS. Regulation of affect by the lateral septum: implications for neuropsychiatry. Brain Res Brain Res Rev. 2004;46:71–117.

10. White SF, Brislin S, Sinclair S, Fowler KA, Pope K, Blair RJR. The relationship between large cavum septum pellucidum and antisocial behavior, callous-unemotional traits and psychopathy in adolescents. J Child Psychol Psychiatry. 2013;54:575–581.

11. Raine A, Lee L, Yang Y, Colletti P. Neurodevelopmental marker for limbic maldevelopment in antisocial personality disorder and psychopathy. Br J Psychiatry. 2010;197:186–192.

12. Crooks D, Anderson NE, Widdows M, Petseva N, Koenigs M, Pluto C, et al. The relationship between cavum septum pellucidum and psychopathic traits in a large forensic sample. Neuropsychologia. 2018;112:95–104.

13. Sarwar M. The septum pellucidum: normal and abnormal. AJNR Am J Neuroradiol. 1989;10:989–1005.

14. Sheehan T, Numan M. The septal region and social behavior. In: Numan R, editor. The behavioral neuroscience of the septal region, New York, NY: Springer New York; 2000. p. 175–209.

15. Shin S, Pribiag H, Lilascharoen V, Knowland D, Wang X-Y, Lim BK. Drd3 Signaling in the Lateral Septum Mediates Early Life Stress-Induced Social Dysfunction. Neuron. 2018;97:195-208.e6.

16. Borelli KG, Blanchard DC, Javier LK, Defensor EB, Brandão ML, Blanchard RJ. Neural correlates of scent marking behavior in C57BL/6J mice: detection and recognition of a social stimulus. Neuroscience. 2009;162:914–923.

17. Bielsky IF, Hu S-B, Ren X, Terwilliger EF, Young LJ. The V1a vasopressin receptor is necessary and sufficient for normal social recognition: a gene replacement study. Neuron. 2005;47:503–513.

18. Lukas M, Bredewold R, Landgraf R, Neumann ID, Veenema AH. Early life stress impairs social recognition due to a blunted response of vasopressin release within the septum of adult male rats. Psychoneuroendocrinology. 2011;36:843–853.

19. Everts HG, Koolhaas JM. Differential modulation of lateral septal vasopressin receptor blockade in spatial learning, social recognition, and anxiety-related behaviors in rats. Behav Brain Res. 1999;99:7–16.

20. Leroy F, Park J, Asok A, Brann DH, Meira T, Boyle LM, et al. A circuit from hippocampal CA2 to lateral septum disinhibits social aggression. Nature. 2018;564:213–218.

21. Wong LC, Wang L, D’Amour JA, Yumita T, Chen G, Yamaguchi T, et al. Effective Modulation of Male Aggression through Lateral Septum to Medial Hypothalamus Projection. Curr Biol. 2016;26:593–604.

22. Hintiryan H, Bowman I, Johnson DL, Korobkova L, Zhu M, Khanjani N, et al. Connectivity characterization of the mouse basolateral amygdalar complex. Nat Commun. 2021;12:2859.

23. Meira T, Leroy F, Buss EW, Oliva A, Park J, Siegelbaum SA. A hippocampal circuit linking dorsal CA2 to ventral CA1 critical for social memory dynamics. Nat Commun. 2018;9:4163.

24. Reddy IA, Pino JA, Weikop P, Osses N, Sørensen G, Bering T, et al. Glucagon-like peptide 1 receptor activation regulates cocaine actions and dopamine homeostasis in the lateral septum by decreasing arachidonic acid levels. Transl Psychiatry. 2016;6:e809.

25. Matthews GA, Tye KM. Neural mechanisms of social homeostasis. Ann N Y Acad Sci. 2019;1457:5–25.

26. Risold PY, Swanson LW. Connections of the rat lateral septal complex. Brain Res Rev. 1997;24:115–195.

27. Maynard KR, Hobbs JW, Phan BN, Gupta A, Rajpurohit S, Williams C, et al. BDNF-TrkB signaling in oxytocin neurons contributes to maternal behavior. ELife. 2018;7.

28. Unable to find information for 3606217.

29. Berton O, McClung CA, Dileone RJ, Krishnan V, Renthal W, Russo SJ, et al. Essential role of BDNF in the mesolimbic dopamine pathway in social defeat stress. Science. 2006;311:864–868.

30. Ito W, Chehab M, Thakur S, Li J, Morozov A. BDNF-restricted knockout mice as an animal model for aggression. Genes Brain Behav. 2011;10:365–374.

31. Huang EJ, Reichardt LF. Neurotrophins: roles in neuronal development and function. Annu Rev Neurosci. 2001;24:677–736.

32. Chao MV. Neurotrophins and their receptors: a convergence point for many signalling pathways. Nat Rev Neurosci. 2003;4:299–309.

33. Song M, Martinowich K, Lee FS. BDNF at the synapse: why location matters. Mol Psychiatry. 2017;22:1370–1375.

34. Rutherford LC, DeWan A, Lauer HM, Turrigiano GG. Brain-derived neurotrophic factor mediates the activity-dependent regulation of inhibition in neocortical cultures. J Neurosci. 1997;17:4527–4535.

35. Marty S, Berzaghi MdaP, Berninger B. Neurotrophins and activity-dependent plasticity of cortical interneurons. Trends Neurosci. 1997;20:198–202.

36. Woo NH, Lu B. Regulation of cortical interneurons by neurotrophins: from development to cognitive disorders. Neuroscientist. 2006;12:43–56.

37. Hill JL, Jimenez DV, Mai Y, Ren M, Hallock HL, Maynard KR, et al. Cortistatin-expressing interneurons require TrkB signaling to suppress neural hyper-excitability. Brain Struct Funct. 2019;224:471–483.

38. Maynard KR, Kardian A, Hill JL, Mai Y, Barry B, Hallock HL, et al. TrkB Signaling Influences Gene Expression in Cortistatin-Expressing Interneurons. ENeuro. 2020;7.

39. Cellerino A, Maffei L, Domenici L. The distribution of brain-derived neurotrophic factor and its receptor trkB in parvalbumin-containing neurons of the rat visual cortex. Eur J Neurosci. 1996;8:1190–1197.

40. Gorba T, Wahle P. Expression of TrkB and TrkC but not BDNF mRNA in neurochemically identified interneurons in rat visual cortex in vivo and in organotypic cultures. Eur J Neurosci. 1999;11:1179–1190.

41. Itami C, Kimura F, Nakamura S. Brain-derived neurotrophic factor regulates the maturation of layer 4 fast-spiking cells after the second postnatal week in the developing barrel cortex. J Neurosci. 2007;27:2241–2252.

42. Huang ZJ, Kirkwood A, Pizzorusso T, Porciatti V, Morales B, Bear MF, et al. BDNF regulates the maturation of inhibition and the critical period of plasticity in mouse visual cortex. Cell. 1999;98:739–755.

43. Zheng K, An JJ, Yang F, Xu W, Xu Z-QD, Wu J, et al. TrkB signaling in parvalbumin-positive interneurons is critical for gamma-band network synchronization in hippocampus. Proc Natl Acad Sci USA. 2011;108:17201–17206.

44. Conner JM, Lauterborn JC, Yan Q, Gall CM, Varon S. Distribution of brain-derived neurotrophic factor (BDNF) protein and mRNA in the normal adult rat CNS: evidence for anterograde axonal transport. J Neurosci. 1997;17:2295–2313.

45. Luo F, Mu Y, Gao C, Xiao Y, Zhou Q, Yang Y, et al. Whole-brain patterns of the presynaptic inputs and axonal projections of BDNF neurons in the paraventricular nucleus. J Genet Genomics. 2019;46:31–40.

46. Fawcett JP, Alonso-Vanegas MA, Morris SJ, Miller FD, Sadikot AF, Murphy RA. Evidence that brain-derived neurotrophic factor from presynaptic nerve terminals regulates the phenotype of calbindin-containing neurons in the lateral septum. J Neurosci. 2000;20:274–282.

47. Ferri SL, Kreibich AS, Torre M, Piccoli CT, Dow H, Pallathra AA, et al. Activation of basolateral amygdala in juvenile C57BL/6J mice during social approach behavior. Neuroscience. 2016;335:184–194.

48. Deng X, Gu L, Sui N, Guo J, Liang J. Parvalbumin interneuron in the ventral hippocampus functions as a discriminator in social memory. Proc Natl Acad Sci USA. 2019;116:16583–16592.

49. Garrido Zinn C, Clairis N, Silva Cavalcante LE, Furini CRG, de Carvalho Myskiw J, Izquierdo I. Major neurotransmitter systems in dorsal hippocampus and basolateral amygdala control social recognition memory. Proc Natl Acad Sci USA. 2016;113:E4914–9.

50. Grishanin RN, Yang H, Liu X, Donohue-Rolfe K, Nune GC, Zang K, et al. Retinal TrkB receptors regulate neural development in the inner, but not outer, retina. Mol Cell Neurosci. 2008;38:431–443.

51. Baydyuk M, Russell T, Liao G-Y, Zang K, An JJ, Reichardt LF, et al. TrkB receptor controls striatal formation by regulating the number of newborn striatal neurons. Proc Natl Acad Sci USA. 2011;108:1669–1674.

52. Burkett JP, Andari E, Johnson ZV, Curry DC, de Waal FBM, Young LJ. Oxytocin-dependent consolation behavior in rodents. Science. 2016;351:375–378.

53. Manvich DF, Stowe TA, Godfrey JR, Weinshenker D. A Method for Psychosocial Stress-Induced Reinstatement of Cocaine Seeking in Rats. Biol Psychiatry. 2016;79:940–946.

54. Beyeler A, Chang C-J, Silvestre M, Lévêque C, Namburi P, Wildes CP, et al. Organization of Valence-Encoding and Projection-Defined Neurons in the Basolateral Amygdala. Cell Rep. 2018;22:905–918.

55. Vrtička P, Sander D, Vuilleumier P. Lateralized interactive social content and valence processing within the human amygdala. Front Hum Neurosci. 2012;6:358.

56. Felix-Ortiz AC, Tye KM. Amygdala inputs to the ventral hippocampus bidirectionally modulate social behavior. J Neurosci. 2014;34:586–595.

57. Beyeler A, Namburi P, Glober GF, Simonnet C, Calhoon GG, Conyers GF, et al. Divergent Routing of Positive and Negative Information from the Amygdala during Memory Retrieval. Neuron. 2016;90:348–361.

58. Oliveira VE de M, Lukas M, Wolf HN, Durante E, Lorenz A, Mayer A-L, et al. Oxytocin and vasopressin within the ventral and dorsal lateral septum modulate aggression in female rats. Nat Commun. 2021;12:2900.

59. Gothard KM. Multidimensional processing in the amygdala. Nat Rev Neurosci. 2020;21:565–575.

60. Rosenberger LA, Eisenegger C, Naef M, Terburg D, Fourie J, Stein DJ, et al. The human basolateral amygdala is indispensable for social experiential learning. Curr Biol. 2019;29:3532-3537.e3.

61. de Gelder B, Terburg D, Morgan B, Hortensius R, Stein DJ, van Honk J. The role of human basolateral amygdala in ambiguous social threat perception. Cortex. 2014;52:28–34.

62. Zheng J, Anderson KL, Leal SL, Shestyuk A, Gulsen G, Mnatsakanyan L, et al. Amygdala-hippocampal dynamics during salient information processing. Nat Commun. 2017;8:14413.

63. Spalletta G, Morris DW, Angelucci F, Rubino IA, Spoletini I, Bria P, et al. BDNF Val66Met polymorphism is associated with aggressive behavior in schizophrenia. Eur Psychiatry. 2010;25:311–313.

64. Ernst C, Marshall CR, Shen Y, Metcalfe K, Rosenfeld J, Hodge JC, et al. Highly penetrant alterations of a critical region including BDNF in human psychopathology and obesity. Arch Gen Psychiatry. 2012;69:1238–1246.

65. Han JC, Thurm A, Golden Williams C, Joseph LA, Zein WM, Brooks BP, et al. Association of brain-derived neurotrophic factor (BDNF) haploinsufficiency with lower adaptive behaviour and reduced cognitive functioning in WAGR/11p13 deletion syndrome. Cortex. 2013;49:2700–2710.

66. Purves KL, Coleman JRI, Meier SM, Rayner C, Davis KAS, Cheesman R, et al. A major role for common genetic variation in anxiety disorders. Mol Psychiatry. 2020;25:3292–3303.

67. van den Berg WE, Lamballais S, Kushner SA. Sex-specific mechanism of social hierarchy in mice. Neuropsychopharmacology. 2015;40:1364–1372.

68. Chovnick A, Yasukawa NJ, Monder H, Christian JJ. Female behavior in populations of mice in the presence and absence of male hierarchy. Aggress Behav. 1987;13:367–375.

69. Li Y, Dulac C. Neural coding of sex-specific social information in the mouse brain. Curr Opin Neurobiol. 2018;53:120–130.

70. Karlsson SA, Haziri K, Hansson E, Kettunen P, Westberg L. Effects of sex and gonadectomy on social investigation and social recognition in mice. BMC Neurosci. 2015;16:83.

71. Elsabbagh M, Divan G, Koh Y-J, Kim YS, Kauchali S, Marcín C, et al. Global prevalence of autism and other pervasive developmental disorders. Autism Res. 2012;5:160–179.

72. Riecher-Rössler A. Sex and gender differences in mental disorders. Lancet Psychiatry. 2017;4:8–9.

73. Boyd A, Van de Velde S, Vilagut G, de Graaf R, O’Neill S, Florescu S, et al. Gender differences in mental disorders and suicidality in Europe: results from a large cross-sectional population-based study. J Affect Disord. 2015;173:245–254.

74. Nievergelt CM, Maihofer AX, Klengel T, Atkinson EG, Chen C-Y, Choi KW, et al. International meta-analysis of PTSD genome-wide association studies identifies sex- and ancestry-specific genetic risk loci. Nat Commun. 2019;10:4558.

75. Seedat S, Scott KM, Angermeyer MC, Berglund P, Bromet EJ, Brugha TS, et al. Cross-national associations between gender and mental disorders in the World Health Organization World Mental Health Surveys. Arch Gen Psychiatry. 2009;66:785–795.

76. Blokland GAM, Grove J, Chen C-Y, Cotsapas C, Tobet S, Handa R, et al. Sex-Dependent Shared and Non-Shared Genetic Architecture Across Mood and Psychotic Disorders. BioRxiv. 2020. August 17, 2020. https://doi.org/10.1101/2020.08.13.249813.

77. Chan CB, Ye K. Sex differences in brain-derived neurotrophic factor signaling and functions. J Neurosci Res. 2017;95:328–335.

78. Puralewski R, Vasilakis G, Seney ML. Sex-related factors influence expression of mood-related genes in the basolateral amygdala differentially depending on age and stress exposure. Biol Sex Differ. 2016;7:50.

79. Carbone DL, Handa RJ. Sex and stress hormone influences on the expression and activity of brain-derived neurotrophic factor. Neuroscience. 2013;239:295–303.

80. Lee G, Gammie SC. GABA(A) receptor signaling in the lateral septum regulates maternal aggression in mice. Behav Neurosci. 2009;123:1169–1177.

81. Nelson RJ, Trainor BC. Neural mechanisms of aggression. Nat Rev Neurosci. 2007;8:536–546.

82. Maynard KR, Tippani M, Takahashi Y, Phan BN, Hyde TM, Jaffe AE, et al. dotdotdot: an automated approach to quantify multiplex single molecule fluorescent in situ hybridization (smFISH) images in complex tissues. Nucleic Acids Res. 2020. May 8, 2020. https://doi.org/10.1093/nar/gkaa312.

83. Schindelin J, Arganda-Carreras I, Frise E, Kaynig V, Longair M, Pietzsch T, et al. Fiji: an open-source platform for biological-image analysis. Nat Methods. 2012;9:676–682.

