## Supplemental Methods and Figures for "The basolateral amygdala to lateral septum circuit is critical for regulating sociability in mice"

**SUPPLEMENTAL INFORMATION

Supplemental Methods**

***Adeno-Associated Viruses***

AAV5-EF1a:EGFP was packaged in serotype AAV5 by the University of Pennsylvania (U Penn) Gene Therapy Vector Core; AAV5-EF1a:Cre was packaged in serotype AAV5 at the University of North Carolina (UNC) Vector Core; AAVrg-EF1a:mCherry-IRES-Cre and AAVrg-EF1a:Flpo were packaged in retrograde serotype by Addgene; AAV8-EF1a:P2A-mCherry-fDIO-Cre and AAV8-EF1a:fDIO-mCherry were packaged in serotype AAV8 by the Stanford University Vector Core. AAV5-EF1a:mCherry-FLEX-dtA was purchased from UNC Vector Core. AAVrg-CB7.CI:EGFP was purchased from UPenn Gene Therapy Vector Core.

***Stereotaxic Surgeries***

Adult male mice (10-12 weeks-old) were deeply anesthetized with a mixture of ketamine/xylazine (ketamine, 100 mg/kg; xylazine, 10 mg/kg; intraperitoneal injection). Stereotaxic surgeries and injections were performed using a stereotaxic apparatus (David Kopf Instruments) and Nanoject III injector (Drummond Scientific), respectively. A craniotomy was performed and adeno-associated virus (AAV) was injected into the target brain region at 2 nL/s through pulled glass micropipette. All injections were done bilaterally. The following coordinates were used: LS, + 0.90 anterior-posterior (AP), ± 0.63 medial-lateral (ML), - 3.45 dorsal-ventral (DV) and + 0.40 AP, ± 0.63 ML, - 3.45 DV; BLA, - 1.55 AP, ± 3.20 ML, - 4.55 DV; ventral CA1 of the hippocampus (vCA1), - 3.05 AP, ± 3.65 ML, - 3.5 DV and - 3.05 AP, ± 2.95 ML, - 4.56 DV. For the virus injection, 140 nL of viral solution per each site was injected into the bilateral LS and BLA and 180 nL of viral solution per each site was injected into the bilateral vCA1. The titers of stock AAV viruses were adjusted to 3.0 x 10^12^ particles/mL before injections. After each viral injection, the glass micropipette was left in place for 10 minutes and then slowly withdrawn. The skin was sutured closed. Mice fully recovered under a heat pad before returning to their home cage.

***Four-trial odor recognition test***

For the 4-trial recognition test with non-social odor cues, we used a three-chamber testing arena with one divider in to create a 2 chamber arena for this test. Mice were habituated to this modified 2 chamber arena for 3 consecutive days (10 min per day) before testing days began. A cotton ball with 5 microliter of natural banana oil (Lorann oils) was put in a perforated 50 mL conical tube with eight 2 mm-diameter holes near the ball, and the scented tube was placed under an inverted metal mesh cup in one chamber. The other chamber contained a mesh cup with nothing in it. We allowed the mice to investigate the two chambers for 2 minutes per trial and completed 3 trials total with a 10 minute break between trials. In a fourth trial, we replaced the scented conical tube with a new one containing a cotton ball with the same volume of natural strawberry oil (Lorann oils). We recorded olfactory investigation of the subject, which was defined as nasal contact with the cup.

***Elevated Plus Maze***

Elevated plus maze (EPM) is used to assess anxiety. The maze (length: 45 cm, width: 10 cm, height from floor: 68 cm) consisted of two enclosed arms and two open arms. Two arms were enclosed by 30-cm high walls and the other two were not. The enclosed arms and the open arms faced each other on opposite sides. A mouse is placed into the intersection of the open and closed arms. Entries to the open arms and time spent in open arms were measured during a 5 min session in a blind manner using a Stopwatch+ program developed by the Center for Behavioral Neuroscience (cbn-atl.org) at Emory University. The maze is cleaned between each animal.

***Fear Conditioning***

Animals were conditioned to associate a 30 s tone (4000 Hz) with a 2 s, 0.6 mA foot-shock. 3 tone-shock combinations were given, with the shock co-terminating with the last 2 s of each tone presentation (“Acquisition” in Figure S1D). Tone-shock combinations were separated by 90 s intervals. 48 h later, mice were placed back into the conditioning chamber to measure recall of the conditioning context (in the absence of the tone) for 180 s (“Retrieval in Figure S1D). Following pre-tone context recall, a total of 23 tones (30 s each) were presented in the absence of the foot-shock for recall of the tone in the fear-associated context (tone/context recall) (“Extinction in Figure S1D). Tones were separated by 5 s intervals. Freezing (cessation of movement) during acquisition and recall sessions was quantified with automated tracking software (FreezeScan; CleverSys, Inc.).

***LS Microdissection***

Adult TrkB^fl/fl^ mice (10 -12 week) received bilateral LS injections with either AAV5-EF1a:EGFP or AAV5-EF1a:Cre. One month after the surgery, mice were anesthetized with isoflurane, brains were removed and immediately sectioned in the coronal plane at 500 μm on a cold metal brain slicer (Zivic instrument), and slices were transferred into ice-cold Hank’s Balanced Salt Solution (HBSS; Gibco). The LS was dissected from brain slices in ice-cold HBSS media, and medial septum was cut out to obtain LS tissues only. These dissected tissues were snap frozen in liquid nitrogen and stored at - 80 °C until further processing for total RNA extraction or western blot analyses.

***RNAScope single-molecule fluorescence In Situ Hybridization***

Brains were extracted and flash-frozen in 2-methylbutane (ThermoFisher) and stored at -80°C until further processing. Coronal sections of the LS (12 μm) across the rostral-caudal axis were taken and mounted onto slides (VWR, SuperFrost Plus). We used the RNAScope Fluorescent Multiplex Kit V1 (Advanced Cell Diagnostics[ACD]). The slides were quickly fixed in 10% buffered formalin at 4°C, washed in 1x PBS, and dehydrated in ethanol. Slides were pre-treated with protease IV solution, and subsequently incubated at 40°C for 2 hours in a HybEZ oven (ACD) with antisense probes for *Ntrk2* (Cat No. 423611) and *Gad1* (Cat No. 400951). Transcript expression was visualized on a Zeiss LSM 700 confocal microscope with a 40x oil-immersion lens. A tiled, z-stacked image of the entire LS from each mouse was acquired in each of the rostral, middle, and caudal positions (refer figure 3A), and images were saved as individual, unstitched z-stacks. Each image was designated as either being part of the dorsal, ventral and intermediate LS subregions, and those from other regions (medial septum, septohippocampal nucleus, etc.) and those that could not be confidently assigned to one subregion were excluded from analysis. These images were analyzed using dotdotdot [[82]](https://sciwheel.com/work/citation?ids=8873960&pre=&suf=&sa=0), a custom MATLAB package for analysis of single molecule fluorescent in situ hybridization. 3D segmentation was performed on all LS images using the CellSegM MATLAB toolbox. From here, DAPI labelled nuclei were isolated and used to measure individual transcripts of *Gad1* and *Ntrk2* within each nucleus. Raw data was exported from MATLAB to a csv and inputted into R to analyze co-localization of transcripts within the nucleus. A threshold of a minimum of 5 transcripts was applied to be included as a positive nuclei for *Gad1* or *Ntrk2*.

***Western Blot Analysis***

Lateral septum (LS) and frontal cortex were microdissected in 1 mm-thick coronal brain sections using a brain slicer matrix (Zivic Instruments) in ice-cold Hanks’ balanced salt solution (HBSS; Gibco) buffer and lysed in lysis buffer [50 mM Tris-HCl, pH 7.6, 150 mM NaCl, 5 mM MgCl_2_, 0.1 % sodium dodecyl sulfate (SDS), 1 mM EDTA, 1% Triton X-100, protease inhibitor mixture (Thermo Fisher)] with brief sonication. Protein concentration was measured with BCA protein assay reagent (Pierce). Equal amounts of proteins (60 ug) were loaded and separated in a 10 % Tris-Glycine gel (Bio-Rad) by SDS-PAGE. Proteins were then transferred to a PVDF membrane (Millipore). The following antibodies were used: anti-TrkB (1:500; 80E3/4603S, Cell Signaling Technologies) antibody and anti-Tubulin (1:20000; DM1A, Sigma–Aldrich) antibody. Quantitative densitometric measurement of immunoblots was performed using the Image J program (NIH) using Tubulin as a loading control.

***Social or Novel Object Exposure for Activity Mapping***

Adult male mice (12-15 weeks old) were individually housed for 1 week before the test. The subjects were exposed to either a social stimulus (novel male mouse) or novel object stimulus (50 mL conical tube) by introducing the stimulus to the home cage of the mouse to be tested. The stimulus was placed in the home cage for 90 s and then removed. After the behavioral stimulus was removed, the mice remained in the home cage for 1.5 h and then killed by transcardial perfusion using PBS and 4 % PFA. Brains were extracted and stored in 4% PFA for 24 h, and cryoprotected in 30% sucrose in 1× PBS/sodium azide (0.05%) for 2-3 d. Brains were then rinsed and stored in PBS at 4 C until further processed.

##### ***Immunohistochemistry***

Coronal sections (50 μm) of the LS were cut on a sliding microtome (Leica) with attached freezing stage (Physitemp), washed in 5% Tween-80 in 1× PBS, and incubated in blocking solution (0.5% Tween-80, 5% normal goat serum in 1× PBS) with agitation for 6–8 h. The sections were then incubated in 1:1000 anti-c-Fos antibody (SySy; cat # 226003) in blocking solution overnight at 4 °C with agitation. The following day, the sections were washed, incubated in 1:1000 goat anti-rabbit AlexaFluor 555 (Sigma) in blocking solution for 2 h with agitation, washed again, and incubated in 1:5000 DAPI (Sigma) in 1× PBS for 20 min. Fos protein fluorescence was visualized on a Zeiss LSM 700 confocal microscope with a 40x oil-immersion lens. Z-stacked image within each LS subregion (dorsal, intermediate, and ventral) was acquired in each of the rostral, middle, and caudal positions. Max projections of individual images were analyzed using FIJI [[83]](https://sciwheel.com/work/citation?ids=24178&pre=&suf=&sa=0) using the 3D object counter. For each experiment, we generated a threshold and voxel number to isolate c-Fos nuclear signal by randomly selecting two images from each condition (four total) and optimized a threshold and voxel number that yielded an accurate representation of c-Fos positive nuclei in each image. Those numbers were applied to the whole data set. Generated csv files from FIJI 3D object counter were compiled and analyzed in R.

**Statistics**

All statistics were performed in RStudio v1.1.463 or Prism v.9.0 (GraphPad Software, Inc.). To determine normality of our datasets we used the Shapiro-Wilk test. Data that was normally distributed was analyzed using two-tailed paired or unpaired *t*-test, adjusting *p*-values for multiple comparisons when necessary. Data that was not normally distributed was analyzed using the Mann-Whitney test. Values are expressed as the means ± s.e.m. Statistical significance was accepted when *P*<0.05.

**Supplemental Figures**
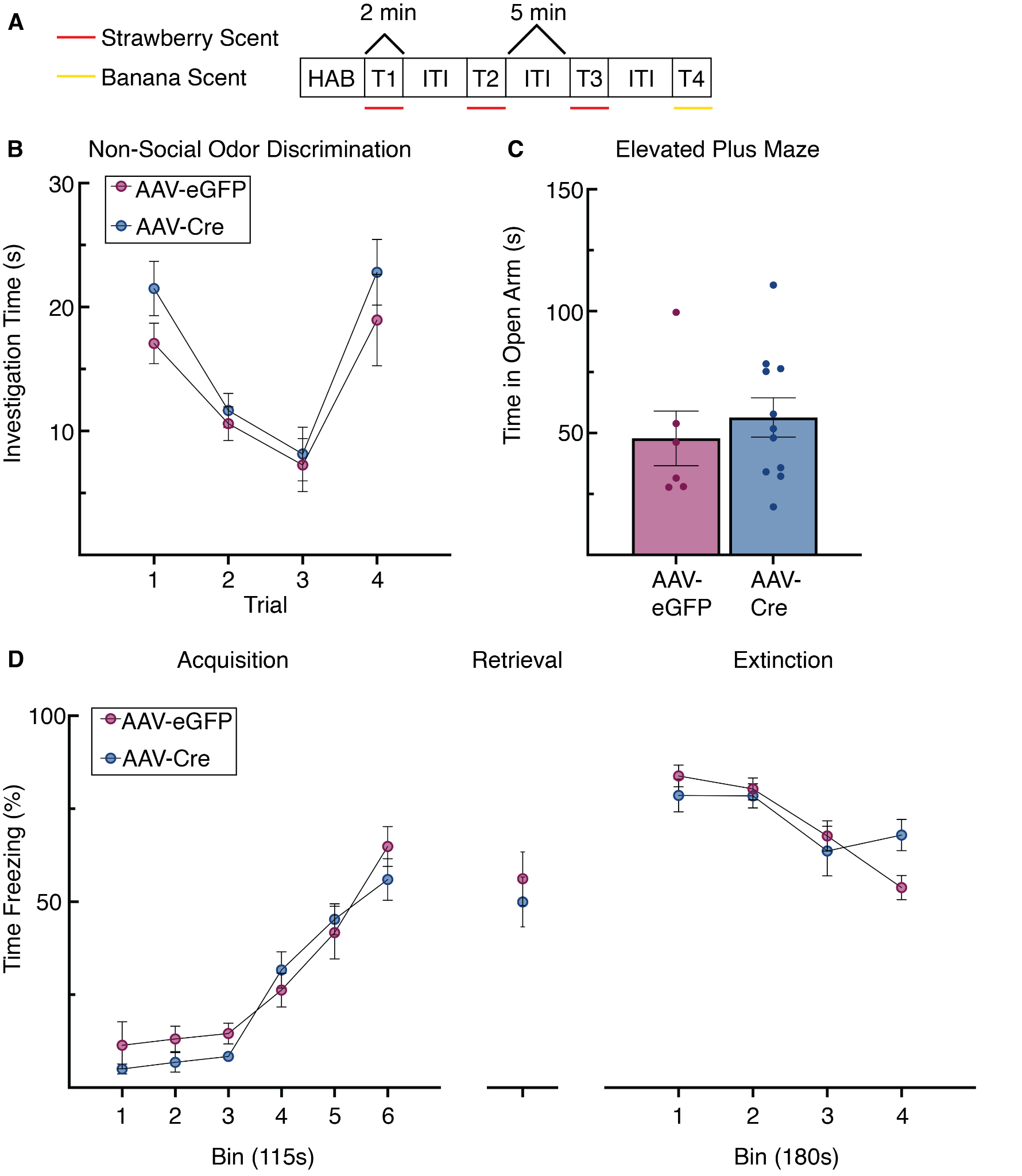

**Supplementary Figure 1.** LS TrkB knockdown does not impair non-social novelty discrimination, fear learning or anxiety. **(A)** Description of the four-trial odor recognition behavioral experiment. **(B)** Knockdown of LS TrkB did not impact ability to discriminate between non-social odors (n=7 AAV-eGFP, n=8 AAV-Cre). Two-way ANOVA revealed no statistically significant interaction between viral group and trial (F(3,18) = 0.276, p = 0.842). **(C)** TrkB Knockdown caused no difference in time spent in the open arm of the elevated plus maze (n=6 AAV-eGFP, n=11 AAV-Cre) (unpaired t-test, t(15) = 0.6266, p=0.5403). **(D)** Acquisition, retrieval and extinction of conditioned fear (n=7 AAV-eGFP, n=11 AAV-Cre). There was no significant difference in acquisition (mixed-factorial analysis of variance, F(5,96) = 1.309, p = 0.2669), retrieval (unpaired t-test, t(16) = 0.612, p = 0.549) and extinction (mixed-factorial analysis of variance, F(3,64) = 1.848, p = 0.1475) of fear conditioning between the groups. Unpaired t-test revealed no significant difference between the groups in retrieval

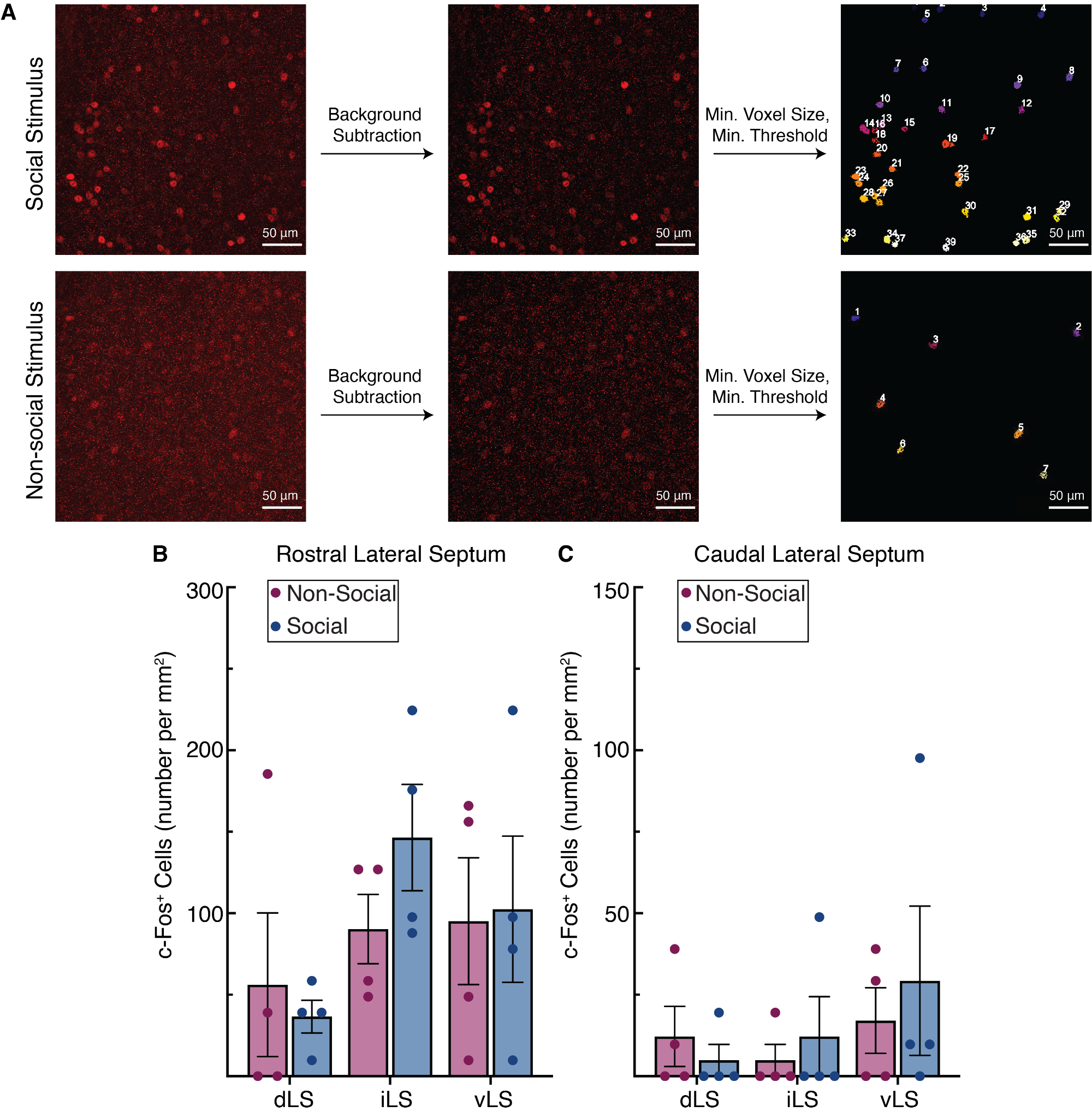

**Supplementary Figure 2.** c-Fos expression across the LS in mice exposed to either a social or non-social stimulus. (A) Schematic illustrating post-processing steps used to quantify c-Fos expression in FIJI. (B) c-Fos induction within the subdivisions of the rostral portion of the lateral septum (C) and the subdivisions of the caudal portion of the lateral septum did not differ between mice exposed to a social and non-social stimulus.

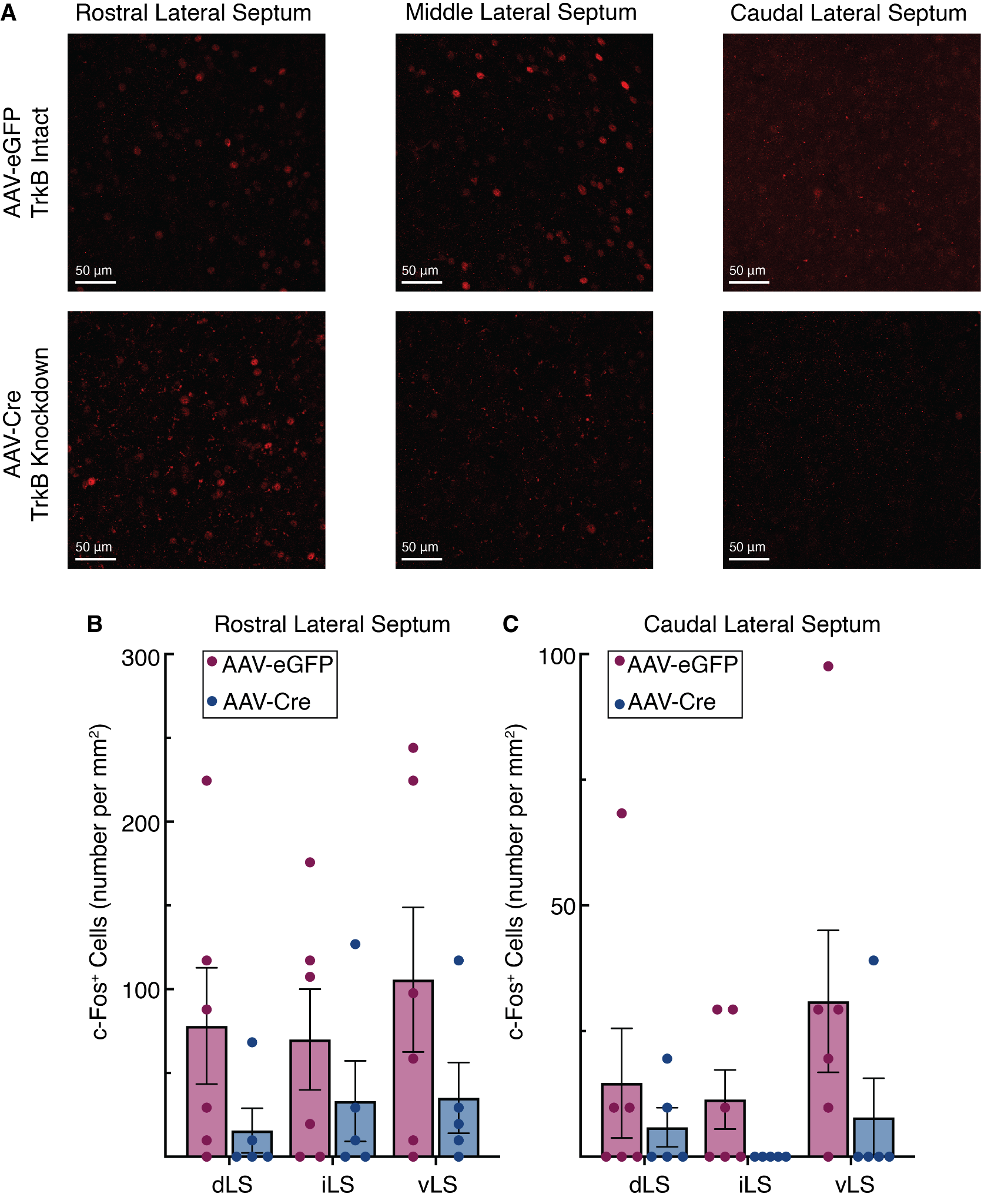

**Supplementary Figure 3.** LS TrkB knockdown does not alter socially induced c-Fos expression in the rostral and caudal LS. (A) Illustration of c-Fos expression in the LS across the rostral caudal divide, representative images are from the iLS. (B) c-Fos induction in response to social stimuli within the subdivisions of the rostral portion of the lateral septum and, (C) the subdivisions of caudal portion of the lateral septum did not differ between mice with knockdown of LS TrkB and mice with intact TrkB expression.
